# Ophthalmate tripeptide is a signaling molecule that regulates striatal neurotransmission

**DOI:** 10.64898/2026.08.03.742629

**Authors:** Dana Shevachman, Archana Proddutur, Shweta Sharma, Peter LeWitt, Gyorgy Lur, Amal Alachkar

## Abstract

Dopamine has long been regarded as the key neurotransmitter governing motor function through the direct and indirect striatal outflow pathways. However, the tripeptide ophthalmate (ophthalmic acid, OA), a glutathione analog, has recently emerged as another regulator of motor function. While the neural mechanisms by which OA regulates motor function remain unknown, we explored how OA behaves as a striatal neuromodulator. Using radiotracer-based uptake and release assays in mouse striatal tissue, combined with whole-cell electrophysiological recordings, we found that OA is taken up through a saturable, glutathione-competitive transport mechanism and is released in a Ca²⁺-dependent manner upon depolarization, consistent with regulated exocytotic release. OA enhanced depolarization-evoked release of γ-aminobutyric acid (GABA) but not glutamate, indicating selective modulation of distinct neurotransmitter systems. OA and dopamine reciprocally regulated one another: OA enhanced dopamine release, while D2 dopamine receptor activation suppressed OA release. Whole-cell recordings from medium spiny neurons (MSNs) showed that OA increased the amplitude of evoked AMPA receptor-mediated currents and shifted short-term plasticity from facilitation toward depression at excitatory synapses onto direct-pathway MSNs (dMSNs). These findings are consistent with an increase in presynaptic glutamate release probability, while sparing synapses onto indirect-pathway MSNs (iMSNs). Together, these findings establish OA as a striatal neuromodulator that interacts with other neurotransmitter systems and preferentially potentiates direct-pathway transmission. This novel discovery identifies a candidate mechanism within the basal ganglia circuitry that governs motor control, with major relevance to Parkinson’s disease and other movement disorders.

## 1. Introduction

Basal ganglia act as a nexus for motor control, procedural learning, and decision making [1–5]. The striatum, the principal input structure of the basal ganglia, integrates glutamatergic inputs from the cortex and thalamus with dopaminergic input from the substantia nigra pars compacta; it also takes in extensive local GABAergic signaling [6]. This integration occurs primarily through the medium spiny neurons (MSNs) that constitute approximately 95% of striatal neurons [7]. Dopamine governs striatal output through two major receptor-defined MSN pathways: D1 dopamine receptor-expressing MSNs of the movement-promoting direct pathway (dMSNs) and D2 dopamine receptor-expressing MSNs of the movement-suppressing indirect pathway (iMSNs) [8–10]. The balance of activity between these two pathways governs basal ganglia output and motor function; disruptions of these pathways or nigrostriatal dopaminergic signaling is central to Parkinson disease (PD) and, possibly, other movement disorders [11]. For more than 6 decades, the independent role of dopamine in the striatum has become a dogma of brain neurophysiology.

Although dopamine is clearly central to striatal motor control, other neurotransmitter and neuromodulator systems and signaling mechanisms also contribute to the regulation of movement [12, 13]. Adding to well-known effects of levodopa, we recently found that this precursor of dopamine also produced a delayed and sustained motor response independently of dopamine synthesis in mouse models of PD; these observation utilized experimental conditions preventing levodopa conversion to dopamine [14]. This prominent motor response coincided with a marked increase in brain levels of ophthalmic acid (ophthalmate, OA) which, in the striatum were elevated approximately 8-fold. Direct intracerebroventricular administration of OA led to dose-dependent rescue of motor deficits in the mouse models of Parkinson’s disease. These findings identify OA as a motor-regulating molecule. However, the mechanisms through which it acts within striatal circuits remained unknown.

Ophthalmic acid is a tripeptide composed of L-γ-Glutamyl-L-2-Aminobutyryl-Glycine (γ-Glu-2AB-Gly). OA is structurally analogous to glutathione (GSH; γ-Glu-Cys-Gly) [15] and was originally [16] isolated from calf lens. This tripeptide has also been detected in various organisms and in tissues that include the mammalian brain [17–22]. OA and GSH share similar biosynthetic pathways. OA is synthesized through the enzymatic actions of glutamate-cysteine ligase (GCL) and glutathione synthase (GS), both of which are involved in GSH production [23]. Studies suggest OA production increases under conditions of oxidative stress in correlation with GSH depletion, thus positioning OA as a potential biomarker of oxidative stress and GSH insufficiency [24–27]. Beyond these observations, research on OA has been limited, and its biological functions remain largely unexplored. However, its recently discovered motor-regulating activity supports the possibility that OA functions as an active neuromodulatory signaling molecule in the brain rather than a passive metabolic byproduct.

Regulated neuromodulatory signaling requires mechanisms for release, action on cellular or synaptic targets, and removal from the extracellular space [28–31]. Whether OA exhibits all of these properties in the striatum has not been established. In particular, it remains unknown whether striatal tissue actively takes up and releases OA, or whether OA modulates established neurotransmitter systems and pathway-specific synaptic transmission.

Here, we tested whether OA functions as a regulated striatal neuromodulator. We combined radiotracer-based uptake and release assays in mouse striatal tissue with whole-cell electrophysiological recordings from identified dMSNs and iMSNs. We found that OA undergoes saturable, GSH-sensitive uptake and calcium-dependent release. OA administration enhanced stimulated dopamine and GABA release, whereas D2 dopamine receptor activation suppressed stimulated OA release. Although OA did not alter bulk glutamate release, it selectively enhanced AMPA receptor-mediated transmission and increased glutamate-release probability at synapses linked to dMSNs. Together, these findings identify OA as a regulated striatal neuromodulator and suggest a pathway-selective synaptic mechanism that may contribute to its motor-promoting effects.

## 2. Materials and Methods

### 2.1. Uptake and Release Assays

#### 2.1.1. Animals

Male Swiss Webster mice (8–10 weeks old) were used for radiolabeled release and uptake studies. For electrophysiological experiments, both male and female C57BL/6J mice were used. All experimental procedures were approved by the Institutional Animal Care and Use Committee (IACUC) of the University of California, Irvine, and were conducted in accordance with NIH guidelines on laboratory animal welfare. Mice were weaned at postnatal day 21 (P21) and group-housed under standard vivarium conditions on a 12 h light/dark cycle, with ad libitum access to food and water.

#### 2.1.2. Brain tissue harvesting and slice preparation for uptake and release assays

Mice were euthanized by CO₂ asphyxiation and brains rapidly removed and transferred to oxygenated (95% O₂/5% CO₂) ice-cold artificial cerebrospinal fluid (aCSF) containing (in mM): 118 NaCl, 4.8 KCl, 2.6 CaCl₂, 1.2 MgSO₄, 25 NaHCO₃, 1.2 KH₂PO₄, 11 glucose, and 0.6 ascorbic acid. Brains were sectioned at 200 µm using a Compresstome (Precisionary Instruments), and two striatal punches were taken from each section using a biopsy punch. All dissection procedures were performed in oxygenated ice-cold aCSF.

#### 2.1.3. Uptake assays

Striatal punches were incubated at 37 °C for 0.5, 1, 5, or 15 min in 150 µL of oxygenated aCSF (pH ∼7.4) containing [³H]-ophthalmate (0.4Ci/mmol, Moravek. Inc.). For competition studies, increasing concentrations of unlabeled glutathione (10⁻¹⁰ to 10⁰ M) were co-incubated with a fixed concentration of [³H]-OA for 5 min. Following incubation, slices were rapidly washed three times with ice-cold aCSF to remove unbound extracellular tracer, then lysed in 1 M NaOH for liquid scintillation counting. For uptake-inhibitor control experiments, slices were pre-incubated for 10 min with nomifensine (10 µM; dopamine transporter inhibitor), nipecotic acid (1 mM; GABA transporter inhibitor), or WAY-213613 (10 µM; EAAT2 inhibitor) prior to and during [³H]-substrate incubation. The time course of [³H]-OA accumulation was fit by nonlinear regression to a one-phase association model in GraphPad Prism (version 11) to estimate the apparent half-time and plateau (expressed as percentage of maximal uptake). GSH competition data were fit to a four-parameter log[inhibitor]-response model to derive the IC₅₀ and Hill slope.

#### 2.1.4. Release assays

For release experiments, three striatal punches were combined per replicate. Slices were preloaded by incubation in [³H]-substrate-containing aCSF for 15 min at 37 °C in the presence of the appropriate uptake inhibitor for the experiment (nomifensine for [³H]-dopamine, nipecotic acid for [³H]-GABA, WAY-213613 for [³H]-glutamate) [32–34]. Following preloading, tissue was washed in aCSF for 30 min at 37 °C to establish a stable baseline, with fractions collected every 5 min. KCl-evoked release was assessed by replacing aCSF with high-KCl aCSF (50 mM KCl, with NaCl reduced isosmotically) for a 5 min stimulation following a 15 min baseline period. Calcium dependence was tested in Ca²⁺-free aCSF containing either 1 mM EDTA or 1 mM EGTA and 2.6 mM MgCl2. For pharmacological experiments, OA (50 µM) or quinpirole (10 µM) was applied 10 min prior to and during KCl stimulation.

Aliquots of perfusate (0.5 mL) were added to 4 mL scintillation fluid and radioactivity quantified using an LS6500 Multi-Purpose Scintillation Counter (Beckman Coulter). Release was expressed as the fractional rate of release (FRR%), calculated as the radioactivity released during a 5 min collection period divided by the total radioactivity remaining in the slice at the start of that period, multiplied by 100.

### 2.2. Whole cell recording

#### 2.2.1. Viral labeling of direct- and indirect-pathway MSNs

To label direct- and indirect-pathway MSNs (dMSNs and iMSNs) for electrophysiology, mice were transduced with cell-type-specific enhancer AAVs driving direct expression of super yellow fluorescent protein 2 (SYFP2) in the dorsal striatum. dMSNs were labeled with a D1-pathway enhancer construct (AiP13044; pAAV-AiE0779m_3xC2-minBG-SYFP2-WPRE3-BGHpA; AAV-PHP.eB; Addgene 191706-PHPeB) while iMSNs were labeled with a D2-pathway enhancer construct (AiP12237; pAAV-AiE0452h-minBG-SYFP2-WPRE3-BGHpA; AAV-PHP.eB; Addgene 191707-PHPeB). Male and female C57BL/6J mice (P24–P35) were anesthetized with isoflurane, and bilaterally injected with the viral construct into the dorsal striatum (coordinates: AP –0.7 mm, ML ±1.5 mm, DV –2.6 mm from bregma; 350 nL per side at 50 nL/min) under aseptic conditions. At least 3–4 weeks was allowed for SYFP2 expression before recording.

#### 2.2.2. Slice electrophysiology

Coronal slices (400 µm) containing the dorsal striatum were prepared on a vibrating microtome (SMZ7000-2, Campden Instruments). Immediately after sectioning, slices were maintained at 32 °C for 15 min in a choline-based cutting solution containing (in mM): 110 choline, 25 NaHCO₃, 1.25 NaH₂PO₄, 3 KCl, 7 MgCl₂, 0.5 CaCl₂, 10 glucose, 11.6 sodium ascorbate, and 3.1 sodium pyruvate, bubbled with 95% O₂/5% CO₂. Slices then recovered for at least 20 min in oxygenated room-temperature recording aCSF containing (in mM): 126 NaCl, 25 NaHCO₃, 10 glucose, 3 KCl, 2 CaCl₂, 1.25 NaH₂PO₄, and 1 MgSO₄ prior to recording.

Recordings were performed near physiological temperature (32–34 °C) in a submersion-type chamber on an Olympus BX51-WI microscope. Whole-cell voltage-clamp recordings targeted fluorescently identified dMSNs or iMSNs in the dorsal striatum, visualized with video-infrared/differential interference contrast microscopy using a 60× water-immersion objective. Glass electrodes (2–4 MΩ) were filled with a cesium-based internal solution for improved space clamp containing (in mM): 135 CsMeSO₃, 10 HEPES, 4 MgCl₂, 4 Na₂ATP, 0.4 NaGTP, and 10 sodium creatine phosphate, adjusted to pH 7.3 with CsOH. Series resistance was 10–20 MΩ. Recordings were made with a MultiClamp 700B amplifier (Molecular Devices), filtered at 4 kHz, digitized at 10 kHz, acquired with WaveSurfer (HHMI Janelia), and analyzed offline in Clampfit 10.7.

AMPA and NMDA receptor-mediated evoked excitatory postsynaptic currents (eEPSCs) were recorded at –70 mV and +40 mV, respectively. A bipolar stimulating electrode was placed approximately 100–150 µm from the recorded neuron in the dorsal striatum and used to evoke paired responses with a stimulus current of 250–500 µA for 500 µs. Inhibitory transmission was blocked with picrotoxin (PTX, 50 µM) and CGP55845 (1 µM) to pharmacologically isolate excitatory responses. In a subset of recordings, the AMPA or NMDA receptor identity of the measured currents was confirmed by bath application of NBQX (10 µM) or AP5 (50 µM), respectively. For each condition, 5–10 sweeps were collected and averaged at a 10 s inter-sweep interval. OA was bath-applied at 10 µM, and paired-pulse and NMDA:AMPA measurements were re-acquired in the same cell after at least 10 min of perfusion.

AMPA receptor currents were measured at –70 mV as the peak amplitude; NMDA receptor currents were measured at +40 mV approximately 50 ms after stimulus onset, a window at which the fast-decaying AMPA component has largely dissipated. The paired-pulse ratio (PPR) was calculated as the second eEPSC amplitude divided by the first (peak 2/peak 1), and the NMDA:AMPA ratio from the averaged peak amplitudes. Recordings were discontinued if series resistance increased by more than 20%, and cells were excluded if lost before or after drug application or if fewer than three sweeps were recorded.

#### 2.2.3. Statistical analysis

Data are presented as mean ± standard error of the mean (SEM) with individual data points where applicable. Analyses were performed in GraphPad Prism (versions 10.4.1–11). The [³H]-OA uptake time course was fit by nonlinear regression to a one-phase association model to estimate apparent half-time and plateau; GSH competition data were fit to a four-parameter log[inhibitor]-response model. For release time courses, two-way ANOVA (treatment x time) was used with Šídák’s multiple comparisons test; for two-condition comparisons, unpaired t-tests were used, with Welch’s correction when standard deviations differed substantially. For paired electrophysiological measurements (aCSF vs OA in the same cell), two-tailed paired t-tests were used. A value of p < 0.05 was considered statistically significant. Sample sizes (n) refer to independent slice preparations for release/uptake experiments and to individual recorded neurons (pairs) for electrophysiology, with animal numbers reported in the figure legends.

## 3. Results

### Ophthalmate is taken up by striatal tissue via a saturable, GSH-competing transporter

To determine whether OA is subject to active cellular uptake in striatal tissue, a prerequisite for transporter-mediated clearance mechanism (Fig. 1A), we first characterized the time course and concentration dependence of [³H]-OA accumulation in mouse striatal punches. [³H]-OA accumulation was time-dependent and approached a plateau by approximately 5 min of incubation, with apparent half-time approximately 0.8 min; plateau approximately 97% of maximal uptake (Fig. 1B,C), consistent with a saturable, carrier-mediated process rather than passive diffusion.

**Figure 1.**
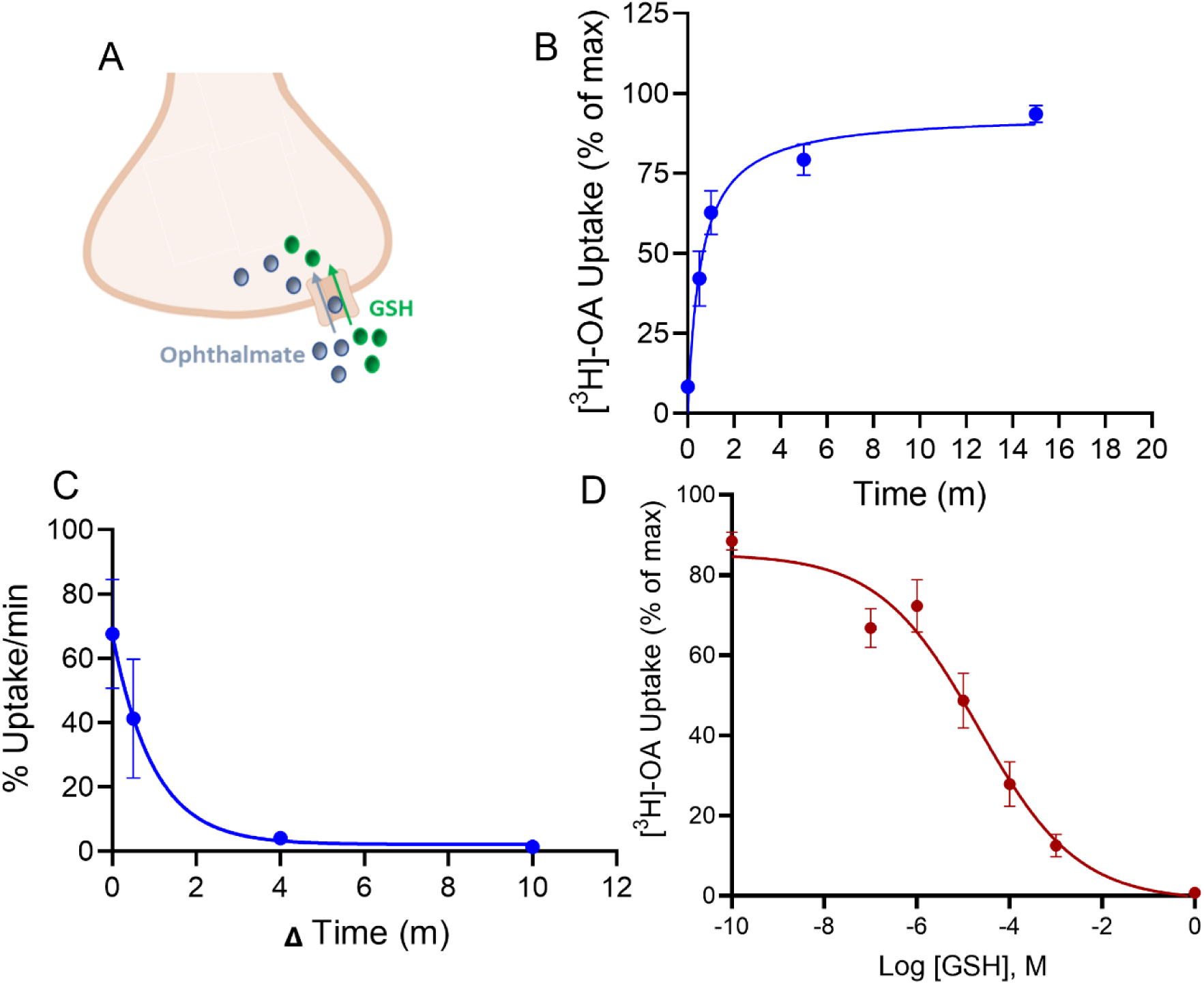
Ophthalmate is taken up by striatal tissue via a saturable, glutathione-competing transport mechanism. **(A)** Schematic of [³H]-OA uptake into a striatal terminal via a transporter shared with glutathione (GSH). **(B)** Time course of [³H]-OA accumulation into mouse striatal punches, fit to a one-phase association model (apparent half-time approximately 0.8 min; plateau approximately 97% of maximal uptake). **(C)** Instantaneous uptake rate (% uptake/min) derived from the time course in (B). **(D)** Concentration-dependent inhibition of [³H]-OA uptake by glutathione (10⁻¹⁰ to 10⁰ M), fit to a four-parameter log[inhibitor]-response model (IC₅₀ = 2.1×10^-5^ M; Hill slope = -0.4050). n = 5 independent experiments, with 3 replicates per condition. Data are presented as mean ± SEM.

To probe the identity of the OA transporter, we tested whether GSH, the structural analog of OA differing only in its central residue, competes for OA uptake. Co-incubation with increasing concentrations of unlabeled GSH (10⁻¹⁰ to 10 M) dose-dependently inhibited [³H]-OA uptake, with an IC₅₀ of 2.1 × 10⁻⁵ M for GSH-mediated inhibition and a Hill slope of −0.4050 (Fig. 1D). This GSH-sensitive uptake indicates that OA and GSH share a common plasma-membrane transport mechanism, providing a candidate route for OA clearance from the extracellular space.

### Depolarization-evoked ophthalmate release is calcium-dependent

A defining property of classical neurotransmitters is Ca²⁺-dependent, depolarization-evoked release from presynaptic terminals. We preloaded striatal slices with [³H]-OA and measured release into the perfusate. Following a stable baseline, application of 50 mM KCl elicited a robust transient increase in [³H]-OA release, with FRR% rising from approximately 15% at baseline to a peak of approximately 44% during stimulation and returning to baseline upon KCl washout (Fig. 2A,B; n = 5, p = 0.0015 KCl vs baseline).

**Figure 2.**
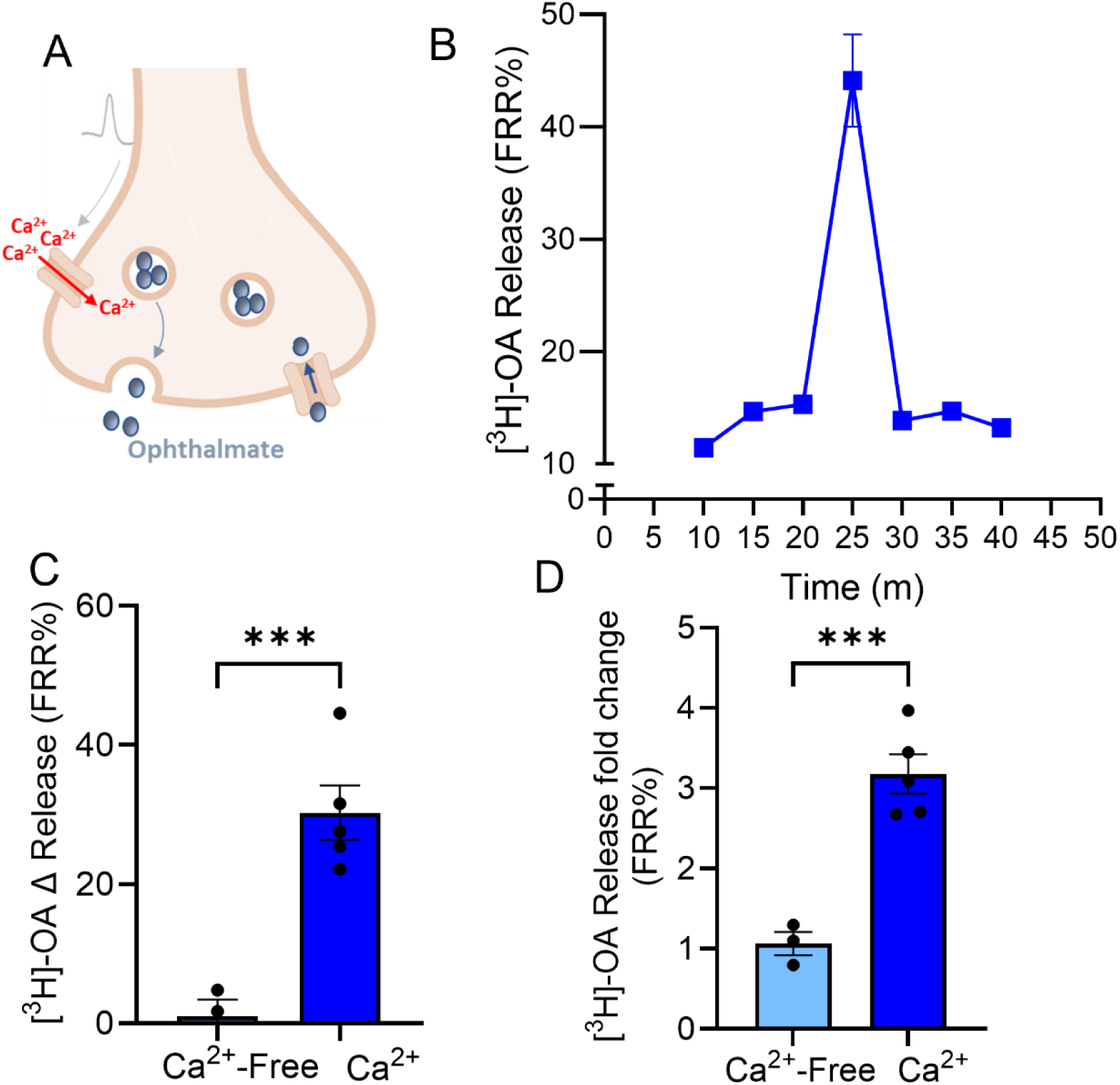
Ophthalmate release is depolarization-evoked and calcium-dependent. **(A)** Schematic of Ca²⁺-dependent [³H]-OA release from a striatal terminal following depolarization. **(B)** Time course of [³H]-OA release (FRR%) with 5 min application of 50 mM KCl at 25 min. **(C)** Change in FRR% during KCl stimulation (ΔRelease) in Ca²⁺-containing vs Ca²⁺-free aCSF; ***p < 0.001. **(D)** Same data as fold change over baseline; ***p < 0.001. n = 4 independent experiments, with 3 replicates per condition. Unpaired Welch’s t-tests. Data are presented as mean ± SEM.

To test the Ca^2+^ dependence of this release, we repeated KCl stimulation in Ca²⁺-free aCSF containing 1 mM EGTA. Under Ca²⁺-free conditions, KCl-evoked [³H]-OA release was abolished: the change in release relative to baseline (ΔFRR%) was reduced from approximately +30% in Ca²⁺-containing aCSF to approximately 0% in Ca²⁺-free aCSF (Fig. 2C; n = 4, p = 0.0007), corresponding to an approximately 3-fold increase over baseline in normal aCSF versus no increase in Ca²⁺-free conditions (p = 0.0003, n = 4; Fig. 2D). The dependence of OA release on extracellular Ca²⁺ is consistent with a vesicular, exocytotic release mechanism analogous to that of classical neurotransmitters.

### Ophthalmate enhances evoked dopamine and GABA release, but not glutamate release

We next asked whether OA modulates the release of classical striatal neurotransmitters. We measured KCl-evoked release of [³H]-dopamine, [³H]-GABA, and [³H]-glutamate in the absence and presence of bath-applied OA (50 µM). To isolate changes in release from changes in reuptake, each experiment was performed in the presence of the relevant transporter inhibitor: nomifensine (10 µM) for dopamine, nipecotic acid (1 mM) for GABA, and WAY-213613 (10 µM) for glutamate. The efficacy of these uptake inhibitors was confirmed in control experiments (Supp Fig. S1).

OA significantly enhanced evoked [³H]-dopamine release, increasing peak FRR% from approximately 105% in vehicle-treated slices to approximately 134% in OA-treated slices (Fig. 3A; n = 4, p = 0.0049), without affecting baseline efflux. Similarly, OA significantly enhanced KCl-evoked [³H]-GABA release, increasing peak FRR% from approximately 98% to approximately 112% (Fig. 3B; n = 4, p =0.0055). In contrast, OA had no significant effect on KCl-evoked [³H]-glutamate release (peak FRR% approximately 46% vehicle vs approximately 42% OA; Fig. 3C; n = 4, p >0.999). The selectivity of OA for dopaminergic and GABAergic but not bulk glutamatergic release argues against a non-specific effect on presynaptic excitability and suggests that OA acts at a defined subset of striatal terminals, consistent with action at a specific receptor.

**Figure 3.**
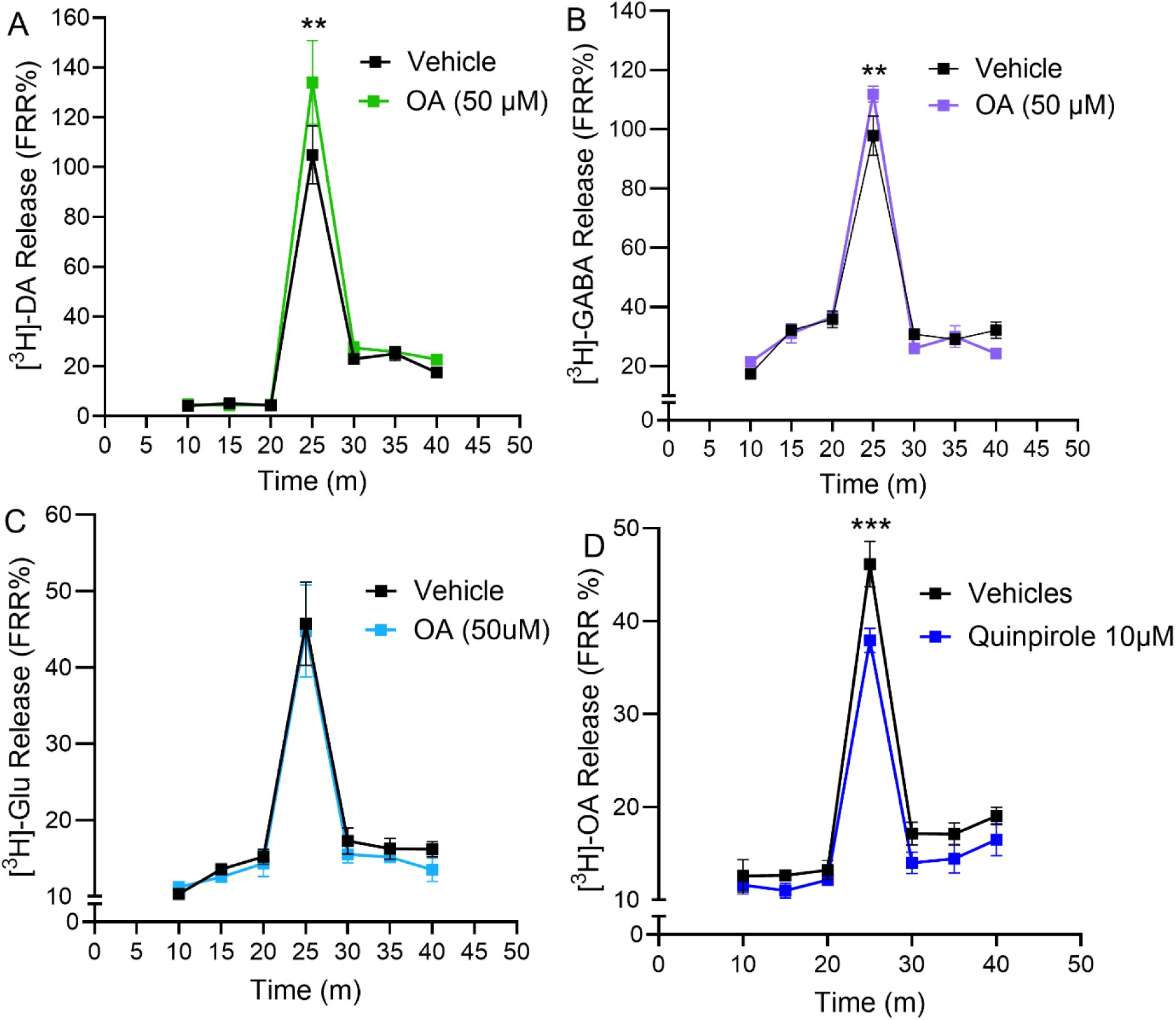
Ophthalmate enhances stimulated dopamine and GABA release, but not glutamate release, and its own release is suppressed by D2 receptor activation. KCl-evoked release of **(A)** [³H]-dopamine, **(B)** [³H]-GABA, and **(C)** [³H]-glutamate in the presence of vehicle or OA (50 µM). Each assay included the appropriate uptake inhibitor (nomifensine, 10 µM, for dopamine; nipecotic acid, 1 mM, for GABA; WAY-213613, 10 µM, for glutamate). OA increased dopamine and GABA release but not glutamate release. **(D)** KCl-evoked [³H]-OA release in the presence of vehicle or the D2 receptor agonist quinpirole (10 µM); quinpirole reduced stimulated OA release. **p < 0.01 (A, B) and ***p < 0.001 (D) at the 25 min peak, two-way ANOVA with Šídák’s multiple comparisons test. n = 4 independent experiments, each with 3 replicates per condition. Data are mean ± SEM.

Our finding that OA enhances dopamine release prompted us to ask whether activation of dopaminergic receptors reciprocally regulates OA release. We tested the D2 dopamine receptor agonist quinpirole because D2 receptors are known to regulate striatal neurotransmitter release. Slices preloaded with [³H]-OA were treated with quinpirole (10 µM) prior to and during KCl stimulation. Quinpirole significantly reduced KCl-evoked [³H]-OA release, decreasing peak FRR% from approximately 46% in vehicle-treated slices to approximately 38% in quinpirole-treated slices (Fig. 3D; n = 4, p = 0.0005). This result demonstrates that D2 receptor activation reduces stimulated OA release.

### Ophthalmate enhances AMPA receptor-mediated excitatory transmission at direct-pathway synapses

To determine how OA acts at individual striatal synapses, we performed whole-cell voltage-clamp recordings from genetically identified dMSNs and iMSNs in acute slices of the dorsal striatum, using cell-type-specific viral enhancers to label each population. AMPA and NMDA receptor-mediated excitatory postsynaptic currents (eEPSCs) were evoked using a paired-pulse protocol, and responses were compared in the same cell before and after bath application of OA (10 µM).

In dMSNs (Fig. 4A), OA significantly increased the amplitude of evoked AMPA responses (aCSF 176.5 ± 39.9 pA, OA 300.9 ± 81.0 pA; p = 0.0348, n = 10, paired t-test, Fig. 4B-D). In addition, OA caused a ∼20% shift in AMPA-mediated short-term plasticity, from facilitation in aCSF to mild depression in OA (AMPA PPR: aCSF 1.14 ± 0.04, OA 0.94 ± 0.02; p = 0.0032, paired t-test, Fig. 4E). In contrast, amplitude and short-term dynamics of NMDA receptor-mediated responses in dMSNs were not impacted by OA (Fig. 4F, G). We also did not detect a significant effect of OA on the NMDA:AMPA ratio (aCSF 2.10 ± 0.38, OA 1.45 ± 0.40; p = 0.166, Fig. 5H). These data indicate that OA has a markedly selective effect on glutamatergic transmission via AMPA receptors in dMSNs.

**Figure 4.**
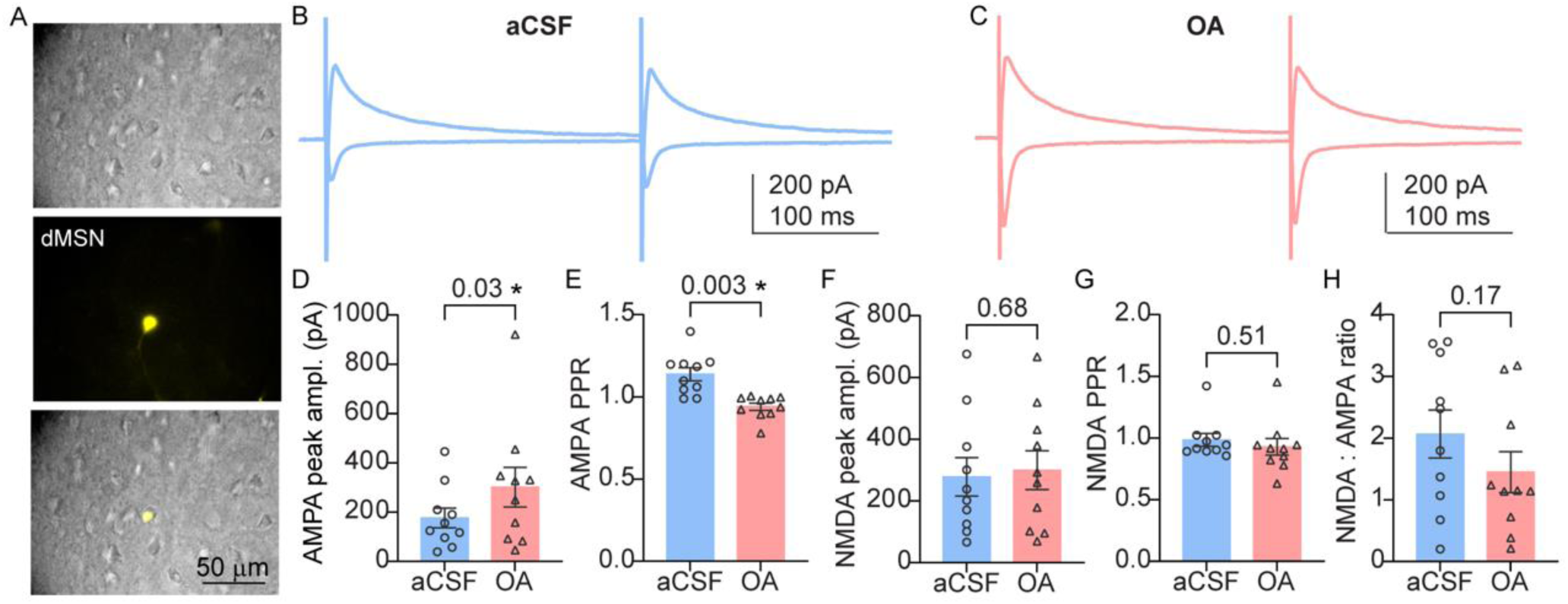
Ophthalmate enhances AMPA receptor-mediated transmission onlo direct-pathway medium spiny neurons (dMSN). **(A)** Top: IR-DIC image of acute striatal slice, middle: fluorescence image of SYFP2-labeled dMSN (D1, AiE0779m_3xC2 enhancer), bottom: merged image. **(B)** Representative traces of whole cell voltage clamp recordings of AMPA (–70 mV) and NMDA (+40 mV) EPSCs evoked by paired-pulse stimulation in aCSF and **(C)** after OA (10 µM) flow in. **(D)** Mean amplitude of AMPA receptor mediated responses. **(E)** Paired pulse ratio of AMPA responses. **(F)** Mean amplitude of NMDA receptor mediated responses. **(G)** Paired pulse ratio of AMPA responses. **(H)** NMDA:AMPA ratio in dMSNs. Bars throughout the figure represent mean ± SEM, individual datapoints are marked as circles (in ACSF, light blue bars) or triangles (following OA flow in, salmon bars). *: p < 0.05, numerical values indicate rounded p-value, paired t-test, n = 10 neurons from 5 mice.

**Figure 5.**
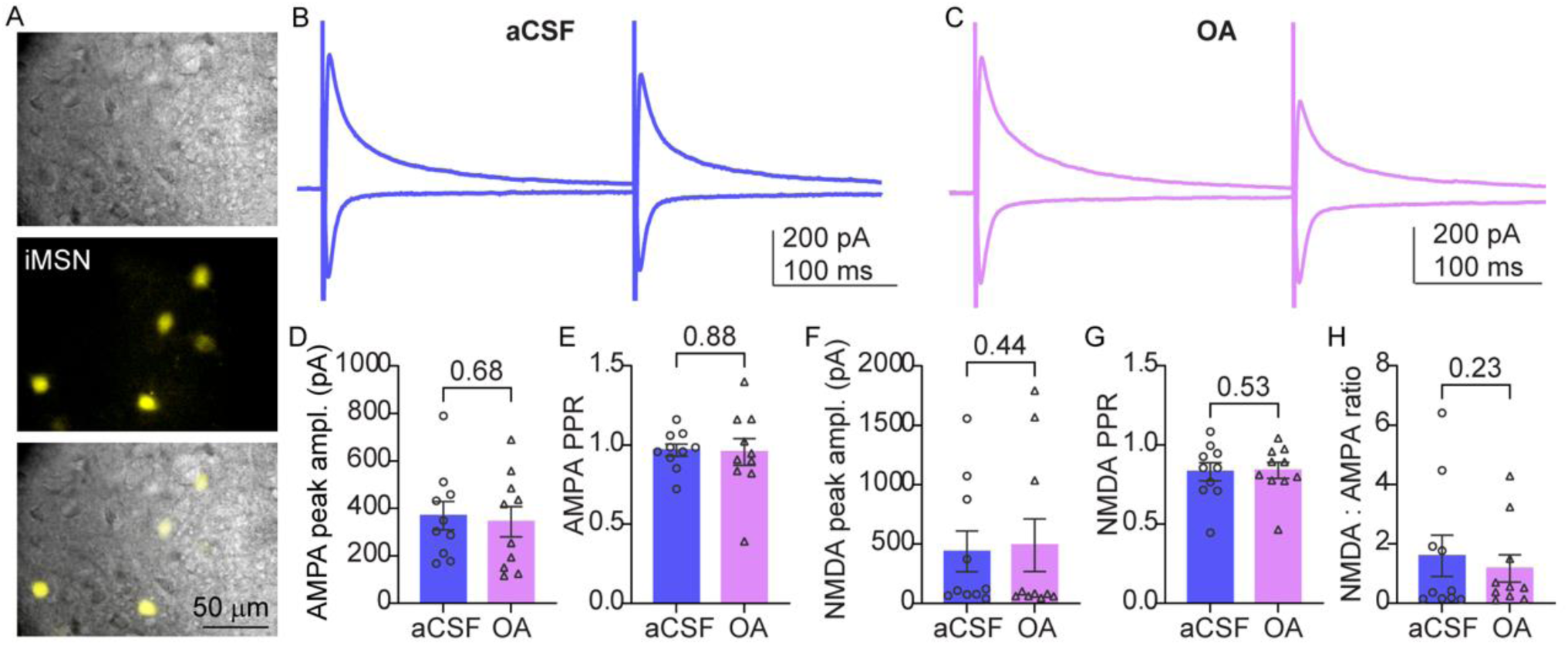
Ophthalmate does not alter excitatory transmission targeting indirect-pathway medium spiny neurons (iMSN). **(A)** Top: IR-DIC image of acute striatal slice, middle: fluorescence image of SYFP2-labeled iMSN (D2, AiE0452h-minBG enhancer), bottom: merged image. **(B)** Representative traces of whole cell voltage clamp recordings of AMPA (–70 mV) and NMDA (+40 mV) EPSCs evoked by paired-pulse stimulation in aCSF and **(C)** after OA (10 µM) flow in. **(D)** Mean amplitude of AMPA receptor mediated responses. **(E)** Paired pulse ratio of AMPA responses. **(F)** Mean amplitude of NMDA receptor mediated responses. **(G)** Paired pulse ratio of AMPA responses. **(H)** NMDA:AMPA ratio in iMSNs. Bars throughout the figure represent mean ± SEM, individual datapoints are marked as circles (in ACSF dark blue bars) or triangles (following OA flow in, magenta bars). *: p < 0.05, numerical values indicate rounded p-value, paired t-test, n = 10 neurons from 5 mice.

Conversely, in iMSNs (Fig. 5A), OA had no significant effect on the amplitude (aCSF 368.5 ± 59.8 pA, OA 343.0 ± 63.8 pA; p = 0.68, n = 10, Fig. 5B-D) or short-term dynamics (aCSF 0.96 ± 0.04, OA 0.95 ± 0.08, p = 0.88, Fig. 5E) of AMPA receptor mediated responses. Similarly to dMSNs, NMDA currents were also unaffected (Fig. 5F, G) and OA had no effect on the NMDA:AMPA ratio (aCSF 1.60 ± 0.68, OA 1.17 ± 0.45; p = 0.226, Fig. 5H). We note that the larger variance of the iMSN NMDA measurements limits the sensitivity of these comparisons; the more tightly distributed AMPA measures, provide stronger evidence for a lack of effect on glutamatergic transmission in iMSNs.

Taken together, OA selectively enhances AMPA receptor-mediated transmission at direct-pathway synapses while leaving indirect-pathway synapses unaffected.

## 4. Discussion

Striatal motor control involves multiple interacting neural circuits and transmitter systems, including dopaminergic, glutamatergic, and GABAergic signaling [12]. Our recent finding that the tripeptide ophthalmate (OA) regulates motor function [14] identified a novel molecule relevant to movement. However, there are lingering questions of how OA acts within striatal circuits, and the nature of its influence on striatal neurotransmission. The present study addresses these questions and shows that OA is actively taken up and released by striatal tissue, modulates dopamine and GABA release, and selectively affects excitatory synapses in the direct (movement-promoting) D1 pathway.

### Ophthalmate exhibits regulated uptake and release by striatal tissue

Our findings establish that OA exhibits several defining features of a regulated neuromodulator in the striatum. First, [³H]-OA is taken up via a saturable, transporter-mediated process that is competitively inhibited by GSH. The latter finding indicates the presence of a clearance mechanism for extracellular OA. This observation aligns with earlier research showing that OA inhibits high-affinity GSH transport (though the latter studies were conducted in liver cells, not brain tissue [35, 36]). The identity of this transporter remains to be established; however, the shared substrate specificity with GSH points to γ-glutamyl-coupled or oligopeptide transport systems. Functional and genetic identification will require further analytical work [37, 38]. Second, depolarization elicits a robust, transient increase in [³H]-OA release that is abolished in Ca²⁺-free medium. This dependence of OA release on extracellular Ca²⁺ is the hallmark of vesicular exocytotic release and it argues against passive efflux or reverse transport as the dominant mechanism [28, 29, 31, 39–44]. Together, these properties (regulated, Ca²⁺-dependent release and a transporter-mediated clearance mechanism) are core features of a neuromodulator. Identifying the receptor that mediates OA’s effects in the striatum is an important next step.

### Bidirectional cross-talk between ophthalmate and dopamine

A central finding of this study is the bidirectional functional interactions between OA and dopaminergic signaling. OA enhanced depolarization-evoked dopamine and GABA release without affecting bulk glutamate release, indicating that OA modulates a subset of striatal terminals rather than acting non-specifically. Conversely, activation of D2 dopamine receptors with quinpirole reduced depolarization-evoked OA release. While this result demonstrates that D2 receptor activation can decrease stimulated OA release, it does not establish the cellular locus of the relevant D2 receptors, which are expressed at multiple pre- and postsynaptic sites in the striatum. This cross-talk may offer a mechanistic explanation for our previously reported observation that brain OA levels rise approximately 20-fold under conditions in which dopamine synthesis is pharmacologically blocked [14]. One possibility is that when dopamine synthesis is reduced, (1) D2 receptor activation falls, (2) the D2-sensitive brake on OA release is relieved, and (3) OA accumulates. Under this model, the OA surge might reflect reduced dopaminergic inhibition of OA release rather than just an incidental metabolic shift. OA may therefore participate in a reciprocal feedback loop with dopamine, and may act as a compensatory signal when dopaminergic tone is low. We note, however, that this interpretation needs to be tested directly: our previous measurements reflect total striatal OA, whereas the present experiments measure stimulated release into the extracellular compartment.

### Ophthalmate enhances excitatory transmission onto direct-pathway MSNs

At the synaptic level, the effect of OA was restricted to dMSNs where it enhanced AMPA receptor mediated responses and reduced the paired-pulse ratio. These findings are consistent with an increase in presynaptic glutamate release probability in response to OA [45, 46]. However, the selectivity of this effect for the AMPA component, with NMDA-mediated responses unchanged, complicates interpretation. An increase in release probability would be expected to enhance both AMPA and NMDA receptor mediated responses. Therefore, an AMPA-only effect would either indicate selectivity to synapses with dominant AMPA receptor content (unlikely given the 1:1 NMDA:AMPA ratio), an NMDA component that is insensitive to changes in release probability, or an additional postsynaptic effect specifically targeting AMPA receptors. Although subtype specific modulation of glutamate receptors has been described previously [47], the present recordings cannot definitively distinguish between the above possibilities.

OA had no detectable effect on any synaptic measures in iMSNs. The cellular basis for this differential effect of OA on glutamatergic transmission targeting dMSNs and iMSNs has not been determined. Aside from differential postsynaptic receptor expression, a possible explanation is that the excitatory afferents targeting dMSNs versus iMSNs arise from molecularly distinct cortical pyramidal populations (e.g. intratelencephalic and pyramidal-tract) [48], which may differ in their sensitivity to OA. One result supporting a differential innervation hypothesis is the difference in baseline short-term dynamics: dMSNs receive facilitating synaptic input while iMSNs innervation is non-facilitating. Establishing a detailed circuit-level explanation would require input-specific manipulation and measurement of pathway output (and which are beyond the scope of this study).

This observed selectivity of OA to selectively modulating excitatory transmission at dMSNs is important because the direct and indirect pathways exert opposing effects on basal ganglia output: direct-pathway activation disinhibits thalamocortical motor circuits and promotes movement, whereas indirect-pathway activation has the opposite effect [3, 7, 49, 50]. A non-selective potentiation of excitatory drive would engage both pathways and produce little net change in motor output. The selective potentiation of dMSN excitatory drive that we observe would be predicted to bias striatal output towards facilitating movement, consistent with OA’s pro-movement behavioral phenotype [14].

The selective effect of OA on AMPA-mediated transmission at dMSN synapses, alongside the absence of any change in bulk KCl-evoked [³H]-glutamate release, is not necessarily contradictory. The bulk release assay samples glutamate from all striatal terminals, as well as non-synaptic and glial sources, whereas the electrophysiological measurements sample excitatory inputs onto identified dMSNs. A change confined to this defined subset of excitatory inputs would be expected to be diluted below detection in a bulk tissue measurement, reconciling the two observations.

### Relationship to motor function and Parkinson disease

Integrating our findings, we propose a working model for the handling and actions of ophthalmate in the striatum (Fig. 6). Our previous work showed that OA administration improves motor function in mouse models of Parkinson’s disease and that brain OA rises under conditions of impaired dopamine synthesis [14]. The present findings show that striatal tissue actively takes up and releases OA and that OA modulates striatal neurotransmission, including release of dopamine and GABA, and excitatory transmission onto dMSNs. These cellular actions provide candidate mechanisms by which OA could influence striatal circuit function. We are cautious, however, about extending these *in vitro* observations into a specific circuit- or disease-level model; the experiments described here were performed on brain slices under defined stimulation conditions, did not manipulate dopaminergic tone, and did not assess motor output. Linking the cellular actions of OA to its behavioral effects will require *in vivo* and input-specific experiments.

**Figure 6.**
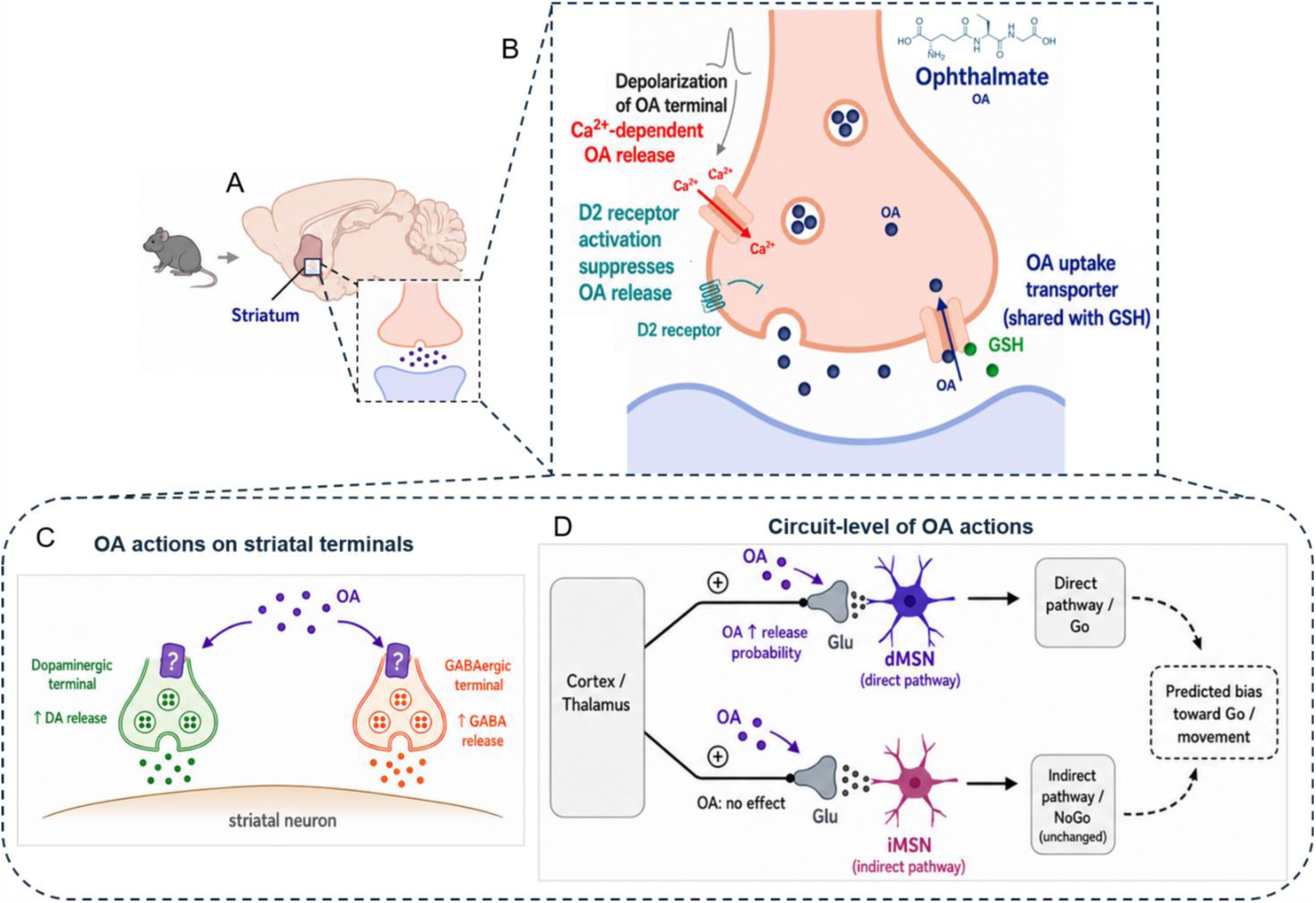
Proposed model of ophthalmate release, uptake, and actions in striatal circuits. Schematic summary of the main findings and their proposed circuit-level implication. **(A)** Anatomical context showing the mouse striatum as the region examined in this study. **(B)** OA release, uptake, and D2-dependent regulation. Depolarization of OA-containing terminals promotes Ca²⁺-dependent OA release, D2 receptor activation suppresses stimulated OA release, and extracellular OA is cleared through a saturable uptake mechanism shared with GSH. **(C)** OA actions on striatal terminals. Released OA enhances dopamine and GABA release from dopaminergic and GABAergic terminals, respectively; the receptor(s) mediating these effects remain unknown. **(D)** Circuit-level actions of OA. At glutamatergic corticostriatal/ thalamostriatal synapses, OA increases release probability onto direct-pathway MSNs (dMSNs), while synapses onto indirect-pathway MSNs (iMSNs) are unchanged. This pathway-selective effect is predicted to bias striatal output toward the direct pathway and movement. Solid arrows indicate experimentally supported findings; dashed arrows indicate inferred circuit-level consequences.

### Limitations and future directions

Several questions remain open for future work. Though Ca²⁺ dependence of [³H]-OA release is consistent with vesicular exocytosis, direct confirmation of vesicular storage will require further study. The receptor mediating OA’s effects in the striatum also remains to be defined; we have previously identified OA as an agonist of the calcium-sensing receptor [14] and whether this receptor underlies the present effects remains to be determined. Finally, the cellular source of striatal OA and the identity of its transporter are not yet known. Together, these define a clear path for future investigation.

## Conclusion

We provide evidence that ophthalmate, a glutathione analog previously regarded as only a passive biomarker of oxidative stress, exhibits the cellular and synaptic properties of a regulated striatal neuromodulator. OA undergoes transporter-mediated uptake, is released in a Ca²⁺- and depolarization- dependent manner, selectively modulates dopaminergic and GABAergic transmission, and selectively increases the amplitude of evoked AMPA responses and release probability at excitatory synapses onto dMSNs, with no detectable effect on iMSNs. OA release is reduced by D2 receptor activation. These findings establish a synaptic and circuit-level mechanism for non-dopaminergic engagement of the direct pathway and expand the neurochemical framework of striatal motor control beyond dopamine.

## Acknowledgements and funding sources

The work of AP and GL was supported by 1R01NS127785.

## Author contributions

DS designed and performed the radiolabeled release and uptake experiments and analyzed the data. AP and GL designed and performed electrophysiological experiments and analyzed the data. AA conceived and supervised the project. DS and AA wrote the manuscript with input from all authors. SS and PL contributed to data analysis and interpretation, and manuscript writing. All authors reviewed and approved the final manuscript.

## AI use statement

Artificial intelligence (AI) tools were used to improve the language and clarity of the manuscript and to generate selected graphical elements incorporated into figures.

## Data availability

All data supporting the findings of this study are available within the article and its supplementary materials. Raw datasets are available from the corresponding author upon reasonable request.

